# Dual roles of a novel oncolytic viral vector-based SARS-CoV-2 vaccine: preventing COVID-19 and treating tumor progression

**DOI:** 10.1101/2021.06.07.447286

**Authors:** Yaping Sun, Wenjuan Dong, Lei Tian, Youliang Rao, Chao Qin, Sierra A. Jaramillo, Erik W. Settles, Shoubao Ma, Jianying Zhang, Kang Yu, Bo Xu, Jiazhuo Yan, Rui Ma, Zhuo Li, Sanjeet S. Dadwal, Bridget M. Barker, Paul S. Keim, Pinghui Feng, Michael A. Caligiuri, Jianhua Yu

## Abstract

The ongoing coronavirus disease 2019 (COVID-19) pandemic is caused by infection with severe acute respiratory syndrome coronavirus 2 (SARS-CoV-2). Cancer patients are usually immunocompromised and thus are particularly susceptible to SARS-CoV-2 infection resulting in COVID-19. Although many vaccines against COVID-19 are being preclinically or clinically tested or approved, none have yet been specifically developed for cancer patients or reported as having potential dual functions to prevent COVID-19 and treat cancer. Here, we confirmed that COVID-19 patients with cancer have low levels of antibodies against the spike (S) protein, a viral surface protein mediating the entry of SARS-CoV-2 into host cells, compared with COVID-19 patients without cancer. We developed an oncolytic herpes simplex virus-1 vector-based vaccine named oncolytic virus (OV)-spike. OV-spike induced abundant anti-S protein neutralization antibodies in both tumor-free and tumor-bearing mice, which inhibit infection of VSV-SARS-CoV-2 and wild-type (WT) live SARS-CoV-2 as well as the B.1.1.7 variant in vitro. In the tumor-bearing mice, OV-spike also inhibited tumor growth, leading to better survival in multiple preclinical tumor models than the untreated control. Furthermore, OV-spike induced anti-tumor immune response and SARS-CoV-2-specific T cell response without causing serious adverse events. Thus, OV-spike is a promising vaccine candidate for both preventing COVID-19 and enhancing the anti-tumor response.

**One Sentence Summary:** A herpes oncolytic viral vector-based vaccine is a promising vaccine with dual roles in preventing COVID-19 and treating tumor progression

## INTRODUCTION

The coronavirus disease 2019 (COVID-19) pandemic, caused by severe acute respiratory syndrome coronavirus 2 (SARS-CoV-2), threatens human health and public safety (*1-3*). Over 3,000,000 people have died worldwide because of SARS-CoV-2 infection by early April 2021 (*4*). Factors such as chronic obstructive pulmonary disease, cardiovascular disease, hypertension, and diabetes mellitus may increase the susceptibility to COVID-19 (*5, 6*). Furthermore, cancer and its treatment usually induce an immunocompromised condition that increases the susceptibility to COVID-19; cancer patients have been reported to have an ∼7-fold higher risk of SARS-CoV-2 infection and an ∼5-fold increased risk of severe COVID-19, as well as an ∼2-fold increased risk of COVID-19 death compared to people without cancer (*7-9*). Sometimes cancer patients even produced few or no antibodies despite having received one of the FDA-approved vaccines, leading them still susceptible to the virus infection (*10*). Therefore, protecting cancer patients from SARS-CoV-2 infection is a high priority for reducing the public health impact of COVID-19 (*11*).

Oncolytic virus (OV), which tends to selectively infect and kill tumor cells but not normal cells, is becoming a promising approach for cancer treatment (*12-14*). OV can activate the immune system against tumor cells by promoting tumor antigen presentation (*15*). Currently, oncolytic herpes simplex virus-1 (oHSV) is one of the most widely used OVs in developing treatment for multiple cancer types (*16*). An oHSV encoding granulocyte–macrophage colony-stimulating factor, named talimogene laherparepvec (T-VEC), is the first and only OV approved by the U.S. Food and Drug Administration (FDA) for cancer treatment (*17, 18*). Many clinical studies have shown that oHSV therapy is relatively safe for cancer treatment (*16, 19, 20*). Most recently, Friedman and his colleagues demonstrated that G207 oHSV is a safe and strong effect on treating pediatric high-grade glioma, with a backbone similar to ours (*20*). Intratumoral administration of oHSV promoted immune cell infiltration into the tumor microenvironment (TME) and activated the immune system (*19-23*). An enhanced immune response can be critical for patients with cancer or COVID-19 or both as all three situations result in an immune-compromised state. oHSV not only stimulates the immune system of cancer patients but also directly lysis tumor cells. Thus, we hypothesize that oHSV vector-based vaccines against SARS-CoV-2 could be a promising approach to protect cancer patients from SARS-CoV-2 infection while initiating, maintaining, or improving anti-tumor immunity.

In this study, we fused the full-length spike (S) protein with oHSV glycoprotein D (gD) to induce S protein expression on the surface of oHSV particles. In order to keep the original features of oHSV, we expressed the transgene encoding the fusion protein at the ICP6 locus driven by the promoter of the HSV-1 immediate early gene IE4/5, while leaving the endogenous gD intact. We show that injection of the OV-spike construct directly induces immune cell activation to produce anti-S-specific antibodies. Long-lasting anti-S neutralization antibodies were produced after injections of OV-spike into tumor-free or tumor-bearing mice. In three different tumor models, melanoma, colon cancer, and ovarian cancer models, OV-spike administration prolonged mouse survival compared to untreated control. Anti-S antibodies induced by OV-spike injection in both tumor-free and tumor-bearing mice inhibited both vesicular stomatitis virus (VSV)-SARS-CoV-2 and live SARS-CoV-2 as well as the B.1.1.7 variant infection. Therefore, OV-spike may have dual roles of preventing cancer progression and serious SARS-CoV-2 infection.

## RESULTS

### COVID-19 patients with cancer have less anti-SARS-CoV-2 immunity than those without cancer

We first compared the levels of anti-S antibodies between COVID-19 patients with and without cancer. The anti-S antibody level was significantly lower in COVID-19 patients with cancer compared to those without cancer (Fig. 1A). Furthermore, we compared the ability of sera from these patients to neutralize the S protein using an in-vitro infection model by VSV-SARS-CoV-2 chimeric virus, which contains an eGFP reporter and is decorated with full-length SARS-CoV-2 S protein in place of the native glycoprotein G (*24*). Compared with the sera from COVID-19 patients without cancer, the sera from those with cancer showed a trend of decreased neutralization against VSV-SARS-CoV-2 infection (Fig. 1B). We also compared the neutralization function of sera from some of these patients against live WT SARS-CoV-2 infection. Similarly, the sera from COVID-19 patients with cancer showed a significantly decreased ability to neutralize live SARS-CoV-2 infection compared to that from COVID-19 patients without cancer (Fig. 1, C and D). These results are consistent with other reports that cancer patients are more susceptible to COVID-19 (*7, 25*).

**Fig. 1.**
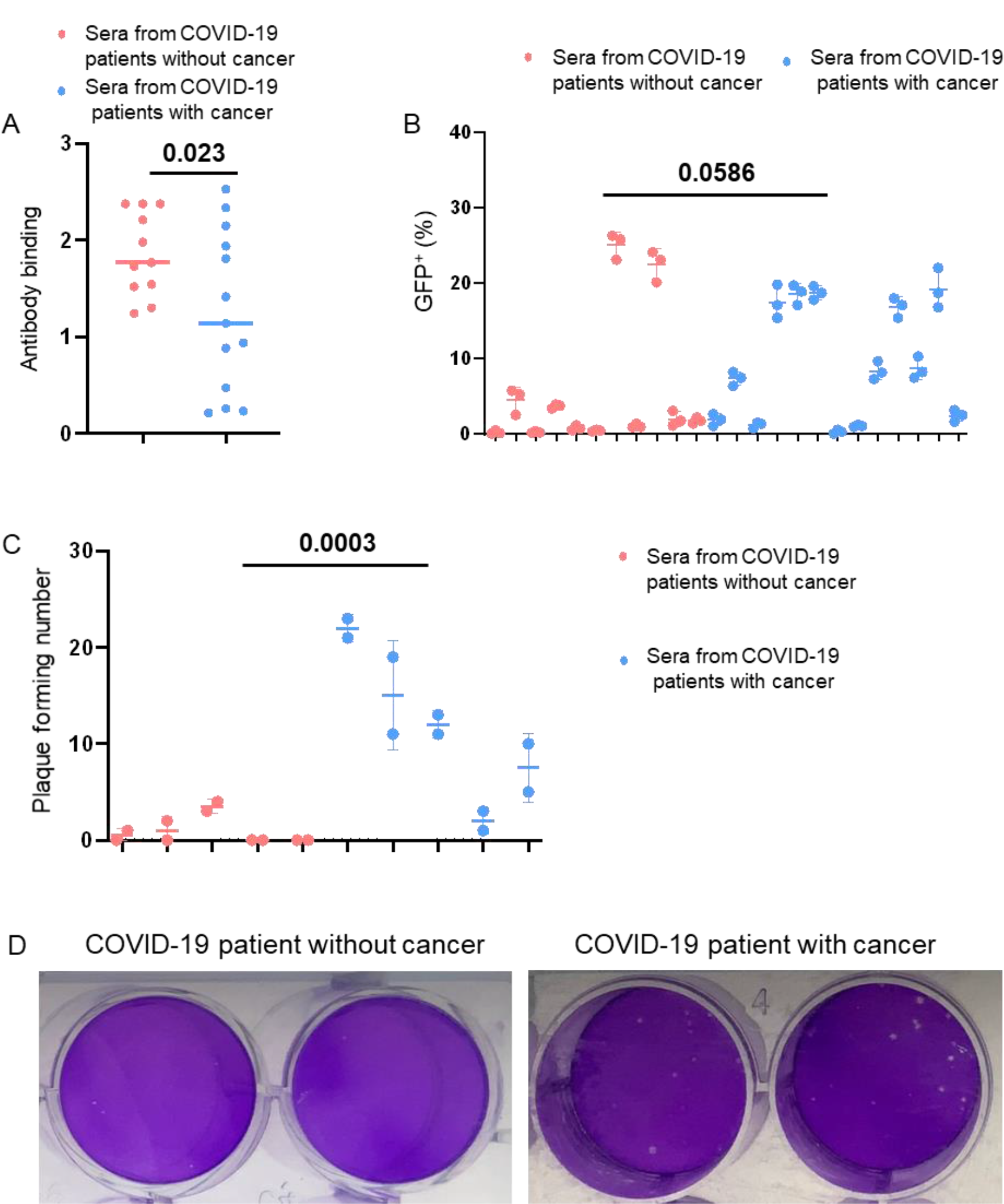
Comparison of antibody production of COVID-19 in patients with and without cancer. **(A)** The presence of anti-SARS-CoV-2 spike (S) protein antibodies in the sera from COVID-19 patients without cancer (n = 11) or with cancer (n = 13) was detected by ELISA using SARS-CoV-2 S protein. **(B)** The ability of serum samples from SARS-CoV-2-infected people with cancer (n = 13) or without cancer (n = 11) to neutralize VSV-SARS-CoV-2 infection in vitro. **(C)** The ability of serum samples from SARS-CoV-2-infected people with or without cancer to neutralize the live SARS-CoV-2 infection in vitro (n = 5 for each group). **(D)** Representative images of the neutralization assay against live SARS-CoV-2 infection of serum samples from COVID-19 patients with or without cancer. Error bars represent standard deviations of triplicates. Two-sample t test for (A), (B) and (C) was applied.

### OV-spike causes SARS-CoV-2 S protein expression on the surface of virus particles and infected cells

Thus, we develop a specialized COVID-19 vaccine with the dual purpose of targeting cancer and boosting anti-SARS-CoV-2 antibody production in cancer patients. The membrane-bound S protein and vector-based vaccines typically work better than non-vector-based vaccines (*26*). We thus fused the full-length S protein with the oHSV gD (S-gD) to induce S protein expression on the surface of oHSV particles (*27-29*) (Fig. 2A). OV-spike was constructed based on the parental oHSV named OV-Q1, which was double-attenuated by inactivating the ribonucleotide reductase gene (ICP6) and deleting both copies of the neurovirulence gene (ICP34.5), thereby limiting its replication to tumor cells and reducing its neurovirulence (*30*). To retain expression of the endogenous gD protein, which is used for oHSV entry into cells, we expressed the transgene encoding the S-gD fusion protein at the ICP6 locus, driven by the promoter of the HSV-1 immediate early gene IE4/5, without interrupting the endogenous gD protein. Thus, OV-spike is designed to maintain endogenous gD expression and ectopically express the SARS-CoV-2 S protein on the surface of viral particles. The genetic maps of WT human HSV-1, OV-Q1, and OV-spike are illustrated in Fig. 2A, and the schematic of OV-spike is shown in Fig. 2B. Negative staining electron microscopy revealed the construct of OV-Q1 and OV-spike (Fig. 2C), and the S protein was verified on the surface of OV-spike but not on the surface of OV-Q1 using immunogold labeling with an anti-SARS-CoV-2 S protein antibody (Fig. 2D). Immunoblot assay of OV-Q1 and OV-spike further confirmed that S protein was expressed in OV-spike viral particles but not the OV-Q1 particles (Fig. 2E). The real-time quantitative PCR results showed that the S protein was highly expressed in the OV-spike-infected cells compared to OV-Q1-infected cells (Fig. 2F). Furthermore, S protein could also be detected on the surface of OV-spike-infected-but not OV-Q1-infected-Vero cells by flow cytometry (Fig. 2G). Collectively, our results indicate that our novel OV-spike vaccine candidate induced S protein expression on the surface of viral particles and the infected cells, therefore, possibly working as a vector-based vaccine.

**Fig. 2.**
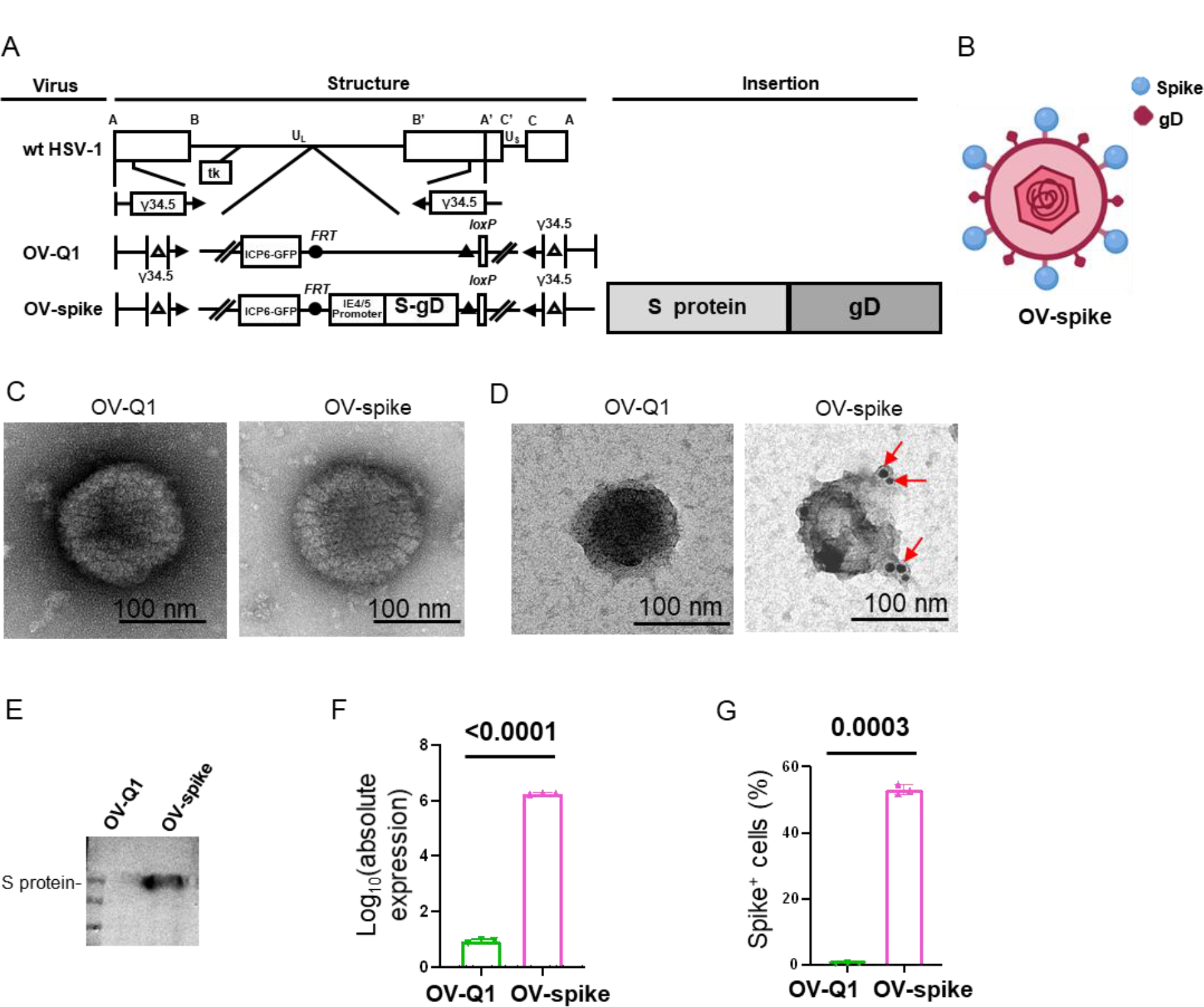
Engineering and validation of OV-spike. **(A)** Schematic map of the oncolytic viruses used in this study. Top: genetic map of wild-type HSV-1. Middle: genetic map of the control oHSV (OV-Q1) with deletion of 2 copies of γ34.5, dysfunctional ICP6, and insertion of the GFP gene. Bottom: genetic map of OV-spike showing the inserted S protein-coding gene fused with glycoprotein D (S-gD). **(B)** The construction model for OV-spike. The blue dots represent S protein fused with the transmembrane and intracellular domain of gD, and the red rhombus shows wild-type gD. **(C)** Transmission electron micrographs of control OV-Q1 (left panel) compared to OV-spike (right panel). **(D)** Immunogold labeling using antibodies against the SARS-CoV-2 S protein for OV-Q1 (left panel) compared to OV-spike (right panel). **(E)** Expression of the SARS-CoV-2 S protein in control OV-Q1 and OV-spike virus particles as determined by immunoblotting assay. **(F)** Expression of the SARS-CoV-2 S protein mRNA in OV-Q1- and OV-spike-infected Vero cells, as measured by quantitative real-time PCR. **(G)** Expression of the SARS-CoV-2 S protein in OV-Q1- and OV-spike-infected Vero cells as detected by flow cytometry. Error bars represent standard deviations of triplicates. Two-sample t test for (F) and (G) was applied.

### OV-spike vaccination induces anti-S antibody production in mouse sera

To test whether OV-spike injection could induce anti-S-specific antibodies, normal BALB/c mice were vaccinated on day 0 with 1×10^6^ or 5×10^5^ plaque-forming units (pfu) of OV-spike via intravenous (i.v.) administration. Mice were administered 1×10^6^ pfu OV-Q1 or saline (mock) as negative controls. On day 14, the mice were boosted with a second dose. Serum samples were collected every 7 days for 10 weeks. High levels of anti-S-specific antibodies could be detected as early as day 21 (Fig. 3A). The production of the antibodies peaked on day 28 and gradually decreased, though a substantial amount still was present on day 70 (Fig. 3B). One hundred percent of tested mice produced anti-S antibodies on days 21 and 28, and over 50% of the mice still produced substantial levels of the antibodies on day 70 (Fig. 3C). Meanwhile, mice of a different strain, C57BL/6, were also i.v. injected with 1×10^6^ pfu of OV-spike to validate the results. The vaccinated C57BL/6 mice produced anti-S-specific antibodies, and the antibody concentration peaked on day 21 post vaccination (fig. S1, A and B). Notably, the anti-S-specific antibodies were present in sera from i.v. vaccinated C57BL/6 mice as early as day 7 (fig. S1, B and C). Furthermore, we also vaccinated C57BL/6 mice intraperitoneally (i.p), with 1×10^6^ pfu or 2×10^6^ pfu OV-spike on days 0 and 14 to test an alternative administration route. Similar results were observed but with a clearer dose-dependent response (Fig. 3, D to F). Most mice started to produce anti-S-specific antibodies rapidly after day 7, and in general, mice in the high OV-spike dose group produced antibodies more quickly than those in the corresponding low dose group (Fig. 3F).

**Fig. 3.**
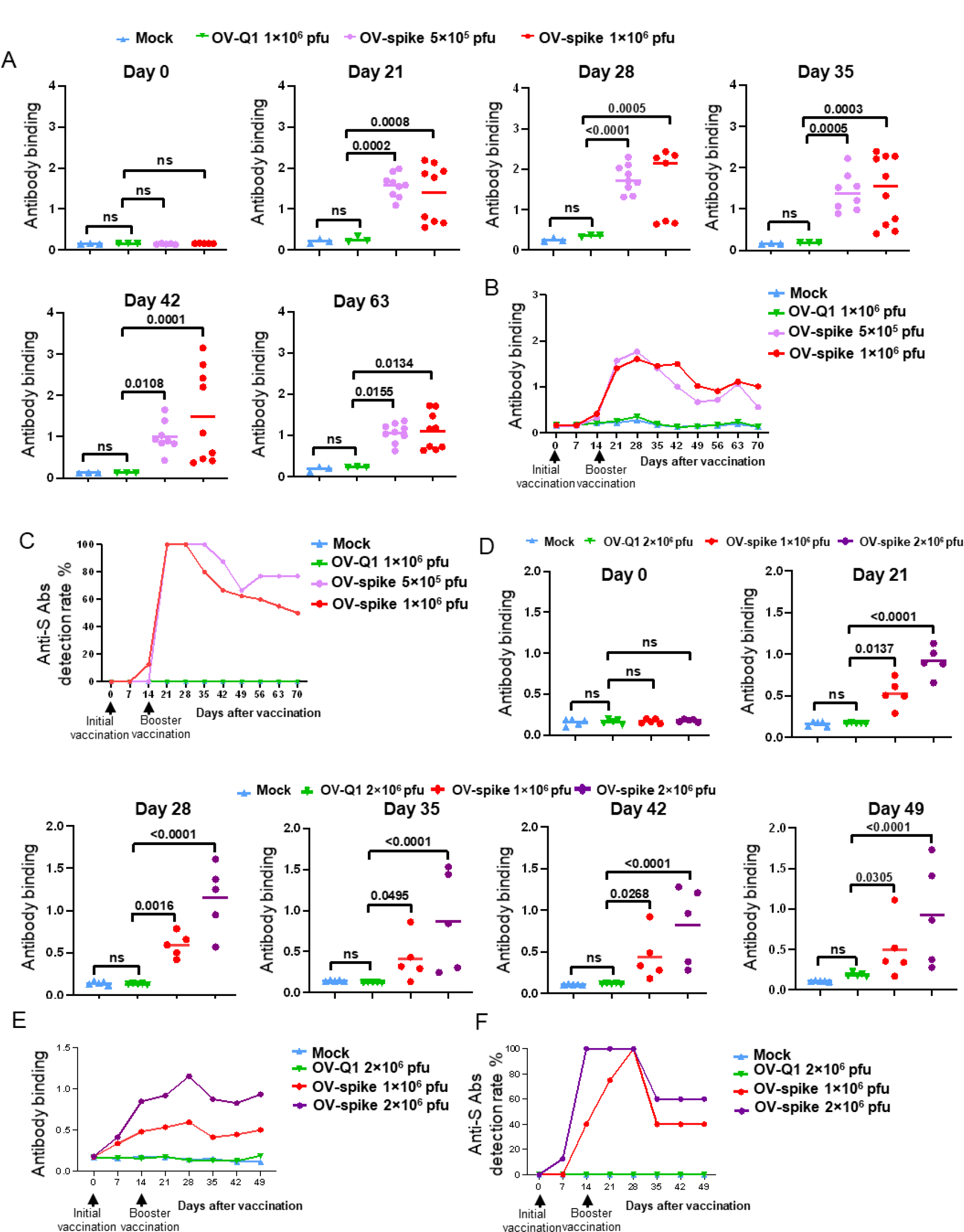
OV-spike vaccination induces anti-S protein production in mouse sera. **(A-C)** BALB/c mice were vaccinated on days 0 and 14 with 1×10^6^ plaque-forming units (pfu) or 5×10^5^ pfu of OV-spike by intravenous (i.v.) administration and 1×10^6^ pfu of OV-Q1 or saline (mock) as a negative control. **(A)** Anti-S protein antibody levels of sera on days 0, 21, 42, and 63, as assessed by an S protein-specific enzyme-linked immunosorbent assay (ELISA). **(B)** Overview of serum anti-S protein antibody levels from days 0 to 70. **(C)** Anti-S antibody production rates in vaccinated mice at the indicated days. **(D-F)** C57BL/6 mice were vaccinated on days 0 and 14 with 1×10^6^ pfu or 2×10^6^ pfu OV-spike by intraperitoneal (i.p.) administration. **(D)** Serum anti-S protein antibody levels on days 0, 21, 28, 35, 42, and 49, as assessed by an S protein-specific ELISA. **(E)** Overview of serum anti-S protein antibody levels from days 0 to 49. **(F)** Anti-S antibody production rates in vaccinated mice at the indicated days. Data in (A) and (D) are shown in mean value, and statistical analyses were performed by one-way ANOVA with P values corrected for multiple comparisons by Bonferroni method multiple comparisons test (n = 3-5 mice for mock group and OV-Q1 group, n = 5-10 mice for OV-spike group).

### Sera from OV-spike-vaccinated mice inhibit infection of VSV-SARS-CoV-2 and wild-type live SARS-CoV-2 as well as the B.1.1.7 variant

We used an enzyme-linked immunosorbent assay (ELISA) to measure the binding affinity of serum samples from vaccinated mice to S protein. Only the sera from OV-spike-immunized mice had a high level of binding affinity with S protein, and the major binding epitope was in S1 subunit with less binding to S2 subunit and nucleocapsid protein (NP) (Fig. 4A). Flow cytometry further confirmed this result, as the sera from OV-spike-immunized mice could efficiently bind to HEK293T cells expressing S protein on the cell surface (Fig. 4B). A neutralization assay with the VSV-SARS-CoV-2 chimeric virus revealed that sera from OV-spike-immunized mice could neutralize viral infection in a dose-dependent manner (Fig. 4C and fig. S2, A and B), and the sera collected from OV-spike immunized mice showed neutralization capacity up to a 320-fold dilution but did not at a 640-fold dilution (Fig. 4, C and D). To further confirm the neutralization function of sera from OV-spike vaccinated mice, prior to infection of Vero cells, live WT SARS-CoV-2 virus was preincubated with the diluted sera from mice immunized with OV-spike or OV-Q1 in a BSL3 lab. The extent of viral infection was determined using an immunoplaque assay. We found that the sera collected from OV-spike-immunized mice significantly reduced live WT SARS-CoV-2 infection in a dose-dependent manner compared to sera from OV-Q1-immunized mice (Fig. 4, E and F). The neutralization capacity of sera from vaccinated mice was also confirmed with a traditional plaque assay (fig. S2C).

**Fig. 4.**
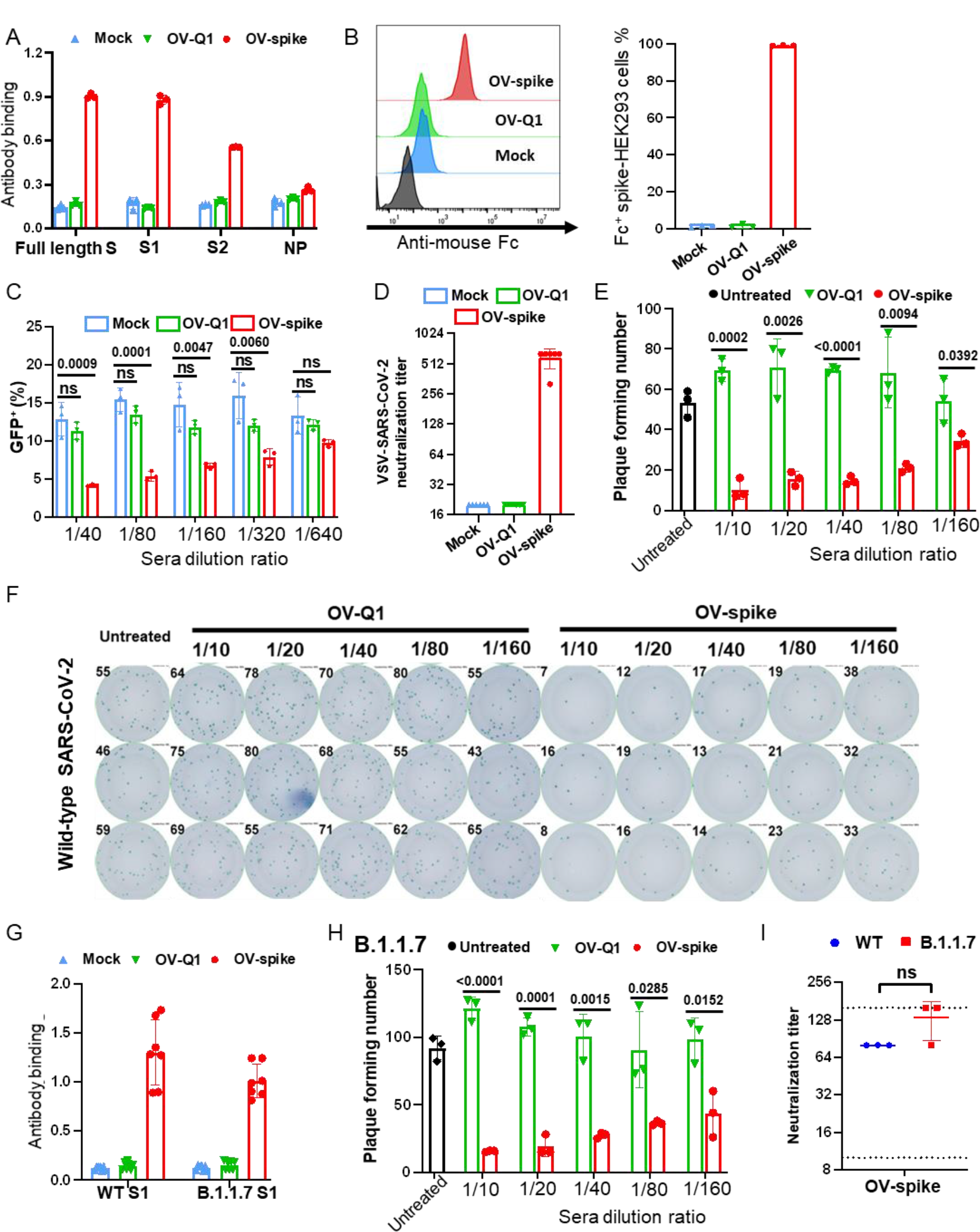
Sera from OV-spike vaccinated mice inhibits both VSV-SARS-CoV-2 and SARS-CoV-2 infection in vitro. **(A)** ELISA-based binding assessment for S protein or S protein subunit 1 (S1) and sera from mice vaccinated with saline (mock), OV-Q1 or OV-spike. **(B)** The left panel shows flow cytometry data for binding between sera from mice vaccinated with saline (mock), OV-Q1 or OV-spike and S protein expressed on 293T cells. The data are summarized in the right panel. **(C)** VSV-SARS-CoV-2 neutralization by sera from mice vaccinated with saline (mock), OV-Q1 or OV-spike at the indicated dilutions. **(D)** The neutralization titer of the sera from vaccinated mice. **(E)** The neutralization against live SARS-CoV-2 virus of the sera from vaccinated mice. **(F)** The image data of the neutralization assay against live SARS-CoV-2 virus. **(G)** The binding assay between S1 protein of wild-type SARS-CoV-2 (WT) or B.1.1.7 variant and sera from mock, OV-Q1, and OV-spike-vaccinated mice was measured by ELISA. **(H)** The neutralization against live B.1.1.7 mutant strain infection of the sera from vaccinated mice. **(I)** The neutralization titer against wild-type strain SARS-CoV-2 (WT) and B.1.1.7 variant of the sera from vaccinated mice. Error bars represent standard deviations of triplicates. Statistical analyses were performed by one-way ANOVA with P values corrected for multiple comparisons by Bonferroni method multiple comparisons test for (C), and two-sample t test with two-tail distribution for (E), (H), and (I).

A variant of SARS-CoV-2, named B.1.1.7 and informally known as the “British variant”, is rapidly spreading internationally. This strain can render SARS-CoV-2 to escape protection from COVID-19 antibodies or existing vaccines (*31*). Therefore, we also tested whether our OV-spike vaccine could prevent infection by the B.1.1.7 variant. First, we compared the binding affinity of S1 subunit from WT and the B.1.1.7 variant to the sera collected from OV-spike-immunized C57BL/6 and BALB/c mice. The binding affinity was similar for the S1 subunits of the B.1.1.7 variant and WT strain (Fig. 4G). We also measured the ability of sera from OV-spike vaccinated mice to neutralize the B.1.1.7 variant. Prior to infection of Vero cells, the live B.1.1.7 variant was pre-treated with the diluted sera from OV-spike or OV-Q1 immunized mice. The immunoplaque assay demonstrated that the sera collected from OV-spike vaccinated mice also showed significant neutralization against infection with the live B.1.1.7 variant in a dose-dependent manner compared to that collected from OV-Q1 groups (Fig. 4H and fig. S2D). Furthermore, mice vaccinated with OV-spike showed no difference of neutralization capacity against live WT strain and the B.1.1.7 variant (Fig. 4I). Together, these data demonstrate that OV-spike vaccination can induce hosts to produce anti-S-specific neutralization antibodies and resist infection by SARS-CoV-2, including the B.1.1.7 variant.

### Vaccination with OV-spike inhibits tumor progression and induces anti-S-specific neutralization antibodies in tumor-bearing mice

To evaluate the in vivo anti-tumor efficacy of OV-spike, we established three mouse tumor models: a melanoma model, a colon adenocarcinoma model, and an ovarian tumor model. For the melanoma mouse model, B16 murine melanoma cells (5×10^5^) were subcutaneously (s.c.) injected into each mouse 5 days before the vaccination, followed by intratumoral injection of OV-spike, OV-Q1, or saline on day 0 and day 2. Tumor progression was monitored by measuring the tumor size. OV-spike and OV-Q1 injection caused similar inhibition of tumor growth in vivo, compared to saline injection (Fig. 5A). Serum samples were collected every 7 days to detect anti-S-specific antibodies. Most of mice in the OV-spike injection group had high levels of anti-S-specific antibodies as early as day 7 (Fig. 5B). We also repeated the experiments with the colon adenocarcinoma tumor model and ovarian tumor model. For the colon adenocarcinoma mouse model, MC38 cells (5×10^5^) were delivered i.p. to each mouse four days before the first dose of OV-Q1 or OV-spike i.p injection (day 0). Another two doses of injections were performed on day 7 and day 14 after the first dose injection. Both OV-spike and OV-Q1 injection increased survival in this model relative to saline injection (Fig. 5C). OV-spike injection also stimulated anti-S-specific antibody production starting on day 7 after vaccination (Fig. 5D). Similar results were obtained from the mouse ovarian tumor model, i.e., OV-spike treatment not only inhibited tumor growth but also produced anti-S-specific antibodies (Fig. 5, E and F, and fig. S3). We also noted that sera from the OV-spike-immunized mice bearing tumors reacted against live WT SARS-CoV-2 and the B.1.1.7 variant; the sera significantly reduced the live WT strain and the B.1.1.7 variant infection compared to OV-Q1-immunized mice bearing tumors (Fig. 5, G and H, and fig. S4). Finally, sera from OV-spike vaccinated mice bearing tumor showed no difference of neutralization capacity against the live WT strain and the B.1.1.7 variant strains (Fig. 5I). There was no significant difference in anti-S-specific antibody production between the tumor models (fig. S5A). The sera from OV-spike-immunized mice with or without tumor showed a similar neutralization function against VSV-SARS-CoV-2 infection (fig. S5B). Thus, our data show that OV-spike can induce anti-S-specific neutralizing antibodies in animals with cancer and that the vaccine has a dual function—restraining tumor progression and inducing anti-S-specific neutralization antibodies.

**Fig. 5.**
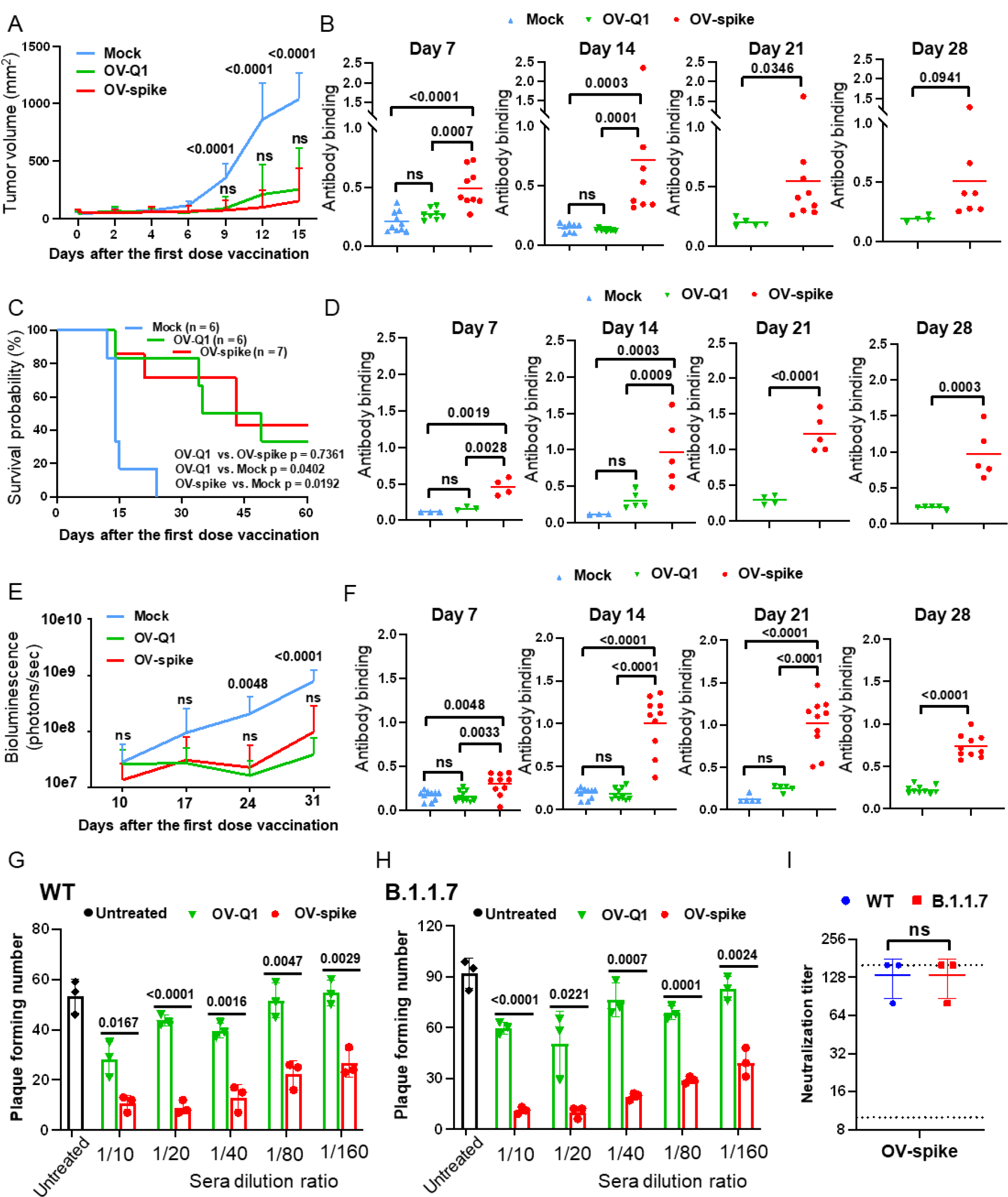
OV-spike vaccine inhibits tumor progression and induces anti-S-specific neutralization antibodies in vivo. **(A and B)** A mouse melanoma tumor model was established by s.c. injection of 5×10^5^ B16 cells. On day 0 and day 2, the mice were intratumorally injected with vehicle control or 1×10^6^ pfu of OV-Q1 or OV-spike. **(A)** Melanoma tumor volume in mice with the indicated treatments. **(B)** ELISA-based assessment of anti-S protein antibody levels in the sera from these mice at the indicated times. **(C and D)** A colon tumor mouse model was established by i.p. injection of 5×10^5^ MC38 cells. On days 0, 7, and 14, the mice were injected i.p. with vehicle control or 2×10^6^ pfu of OV-Q1 or OV-spike. **(C)** Survival of mice with the indicated treatments. **(D)** ELISA-based assessment of anti-S protein antibody levels in the sera from these mice at the indicated times. **(E and F)** A mouse ovarian tumor model was established by i.p. injection of 1×10^6^ ID8 cells. On days 0, 7, and 14, the mice were injected i.p. with vehicle control or 2×10^6^ pfu of OV-Q1 or OV-spike. **(E)** The tumor volume of mice with the indicated treatments. **(F)** ELISA-based assessment of anti-S protein antibody levels in the sera from these mice at the indicated times. **(G and H)** The neutralization against live wild-type SARS-CoV-2 strain (WT) (G) and B.1.1.7 variant (H) infection of the sera from vaccinated mice bearing tumors. **(I)** The neutralization titer against live wild-type strain (WT) and B.1.1.7 variant infection of the sera from vaccinated mice bearing tumors. Error bars represent standard deviations. Data in B, D, and F are shown in mean value. Statistical analyses were performed by one-way ANOVA with P values corrected for multiple comparisons by Bonferroni method multiple comparisons test for (A), (B), (D), and (F) (n = 3 to 10). Two-sample t test with two-tail distribution was applied for (G), (H) and (I).

### Vaccination with OV-spike activates cellular immune responses in both tumor-free and tumor-bearing mice

To evaluate immune system activation after OV-spike vaccination, tumor-free BALB/c mice were injected with OV-spike on days 0 and 14. SARS-CoV-2 specific T cells were analyzed after ex vivo antigen stimulation with an S peptide mixture. An enzyme-linked immunospot (ELISpot) assay showed that the S peptide mixture stimulated significantly more splenic cells to produce interferon gamma (IFNγ) in the OV-spike-vaccinated group than the OV-Q-and saline-vaccinated groups (Fig. 6A and fig. S6A). Consistent with these results, a flow cytometric analysis following ex vivo antigen stimulation using an S peptide mixture showed that OV-spike vaccination significantly increased the percentage of IFNγ^+^ S-specific CD4^+^ and CD8^+^ T cells (Fig. 6, B and C) compared to vaccination with OV-Q1 or saline. However, OV-spike vaccination did not change the percentage of natural killer (NK) cells and NK cell activation (Fig. 6, D and E). In our mouse ovarian tumor model, OV-spike vaccination also induced more IFNγ-producing cells after ex vivo antigen stimulation using an S peptide mixture than vaccination with OV-Q1 or saline, suggesting that OV-spike can also induce antigen-specific T cells in tumor-bearing mice (Fig. 6, F to H, and fig. S6B). Similar to the results with tumor-free mice, there was no difference in the number of NK cells among the saline-, OV-Q1-, and OV-spike-vaccinated mice (Fig. 6I); however, unlike in the tumor-free mice, the percentage of activated NK cells was greater in the OV-Q1- and OV-spike-vaccinated groups than the saline group (Fig. 6J). These ex vivo results are consistent with those collected from in vivo mouse tumor models, wherein OV-spike stimulates anti-S-specific neutralizing antibodies, but both OV-spike and OV-Q1 have anti-tumor effects.

**Fig. 6.**
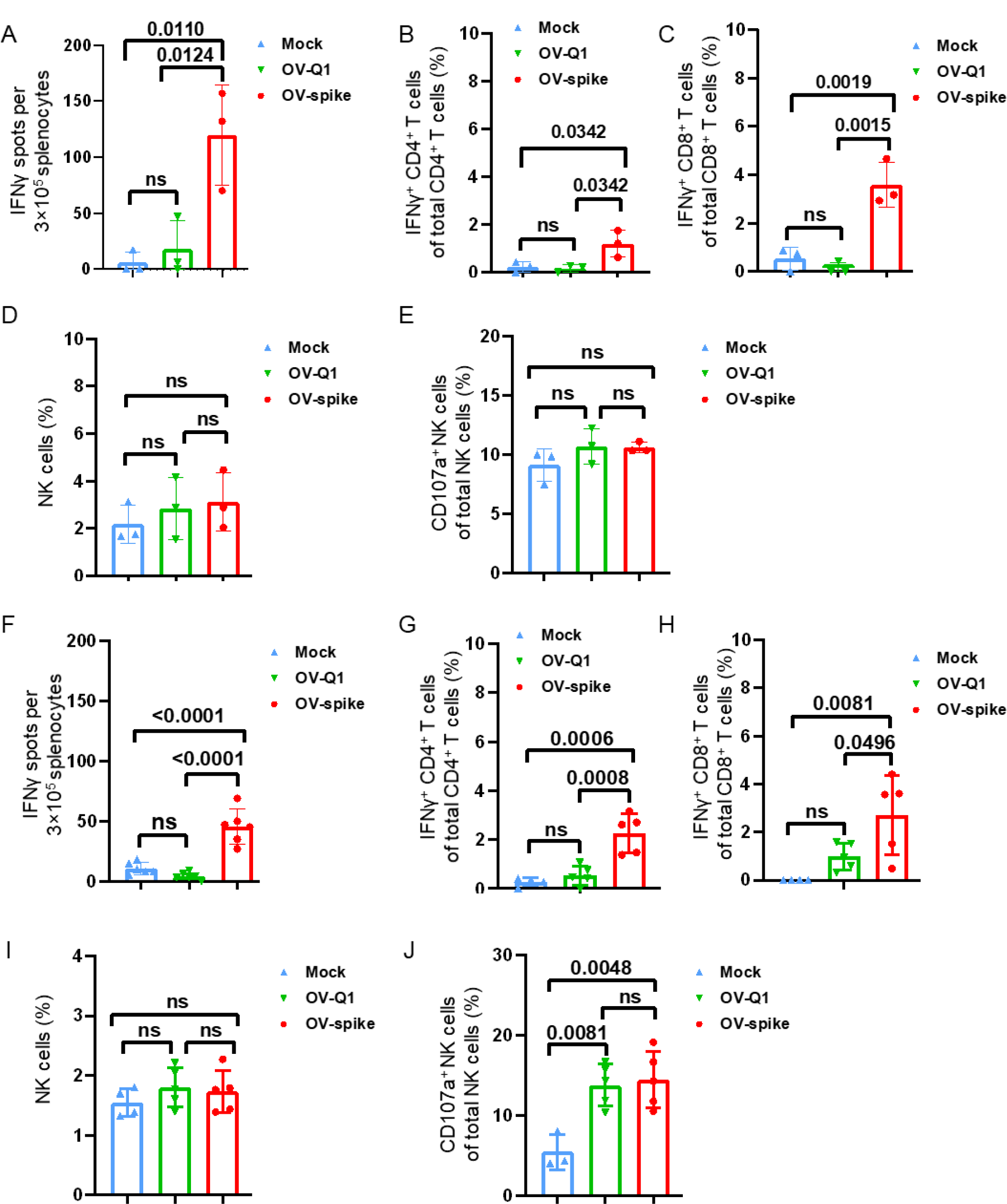
OV-spike vaccine activates immune responses in mice. **(A)** Cellular immune responses of splenocytes as assessed using interferon gamma (IFNγ) ELISpot assays in vaccinated non-tumor-bearing BALB/c mice. **(B and C)** The percentage of IFNγ+CD4^+^ and IFNγ^+^CD8^+^ T cells after exposure to pooled S peptides from splenocytes extracted from mice vaccinated with saline (mock), OV-Q1, or OV-spike, as analyzed by flow cytometry. **(D and E)** The percentage of natural killer (NK) cells and activated (CD107a^+^) NK cells from splenocytes extracted from mice vaccinated with saline (mock), OV-Q1, or OV-spike, as analyzed by flow cytometry. **(F)** Cellular immune responses of splenocytes as assessed using IFNγ ELISpot assays in vaccinated mice bearing ID8 tumors after ex vivo antigen stimulation using an S peptide mixture. **(G and H)** The percentage of IFNγ^+^CD4^+^ and IFNγ^+^CD8^+^ T cells after exposure to pooled S peptides from splenocytes extracted from ID8 tumor-bearing mice vaccinated with saline (mock), OV-Q1, or OV-spike, as analyzed by flow cytometry. **(I and J)** The percentage of NK cells and activated (CD107a^+^) NK cells from splenocytes extracted from ID8 tumor-bearing mice vaccinated with saline (mock), OV-Q1, or OV-spike, as analyzed by flow cytometry. Error bars represent standard deviations, and statistical analyses were performed by one-way ANOVA with P values corrected for multiple comparisons by Bonferroni method multiple comparisons test (n = 3 to 6).

### Lack of side effects observed after OV-spike vaccination

Vaccine safety is a vital concern in clinical application. We thus evaluated the adverse effects of OV-spike vaccine. The same dose of OV-Q1 or OV-spike was injected into tumor-free mice on day 0. We observed that the injection sites had no significant redness and swelling (data not shown). We measured mouse temperature and body weight at the indicated timepoints. Neither fever nor weight loss was observed after immunization (fig. S7, A and B). Histological analysis of the different organs, including the lung, brain, liver, and kidney, showed no substantially notable changes in the OV-spike-vaccinated group compared to the saline and OV-Q1 groups (fig. S7C). These preclinical results indicate a favorable safety profile for OV-spike.

## DISCUSSION

Although there are many vaccines against SARS-CoV-2 in various stages of development and distribution (*32-34*), none are tailored to the needs of people with cancer. To generate OV-spike, a dual functional SARS-CoV-2 vaccine that also halts tumor progression, we modified the oncolytic virus oHSV to express S protein on its surface. Vaccination with OV-spike induced anti-S neutralization antibody production in both BALB/c and C57BL/6 mice. Furthermore, OV-spike not only reduced tumor growth in mice but also prevented SARS-CoV-2 infection of both the wild-type and the B.1.1.7 variant, whereas the parental OV, OV-Q1, only reduced tumor growth. OV-spike vaccination did not cause obvious or severe adverse effects in mice. These results indicate that OV-spike could be a safe and effective dual-functional vaccine to elicit an anti-tumor response while providing protection from SARS-CoV-2 infection to cancer patients.

Cancer patients are relatively immunocompromised with impaired immune cell function and therefore more susceptible to SARS-CoV-2 infection with a poorer response to vaccination (*5, 10, 35*). As such, COVID-19 can cause more severe symptoms in cancer patients. Many reports show the morbidity and mortality rates of cancer patients with COVID-19 are much higher when compared to those of patients without cancer (*7, 8, 25*). Therefore, effective and specific vaccines stimulating stronger immune responses should be urgently developed to protect cancer patients from SARS-CoV-2infection. Most anti-SARS-CoV-2 vaccines attempt to induce anti-viral immune responses against multiple viral proteins, including the S protein (*36-38*). They are either non-vector-based (e.g., RNA-based) (*39*) or vector-based vaccines (e.g., adenovirus-based) (*40*). However, our OV-spike is a unique vector-based vaccine candidate, as it can also have an oncolytic function, rendering it be more suitable for cancer patients. Compared to other vaccines, our OV-spike vaccine not only can produce both anti-tumor and anti-virus immune responses but also may provoke stronger immune responses than other vaccines because of at least two unique features. First, the vaccine is designed such that OV-spike specifically infects tumor cells and induces the tumor cells to express S proteins on the tumor cell surface or release the S antigens after oncolysis. This can amplify the anti-SARS-CoV-2 immune response. Second, after the tumor cells are lysed by OV-spike, they produce tumor-specific antigens that can induce anti-tumor immunity. The anti-viral and anti-tumor immune responses may cross-react with each other to destroy both tumor cells and SARS-CoV-2 virus particles, as the stimulated adaptive immune response can boost the immune responses of innate immune cells (*41*), which may not distinguish tumor killing from virus clearance or vice versa. The amplified immune responses could be beneficial for people with cancer, who are usually immunocompromised and/or have lymphopenia, which can become more extreme after SARS-CoV-2 infection.

Our current study provides proof-of-concept for dual roles of OV-spike in multiple tumor models. In clinical practice, we anticipate this vaccine could be best suited for local injection into solid tumors, such as melanoma and sarcoma. As the vaccine can specifically infect and be amplified by tumor cells and induce the immune amplification mentioned above, the vaccine may be amenable to low-dose injection, especially for people with early-stage cancer, which can lessen the risk of side effects. Although our vaccine candidate fits cancer patients, we cannot exclude the possibility to use the vaccine for non-cancer patients as well because the immune cells can respond to the S protein that we designed and proved to express on the surface of viral particles. Also, considering the observed strong immune response to OV-spike and the immunocompromised or lymphopenic condition of people with severe COVID-19 at the late stage (*42*), OV-spike may also serve as a therapeutic agent not only for cancer but also for COVID-19. Therefore, our vaccine can have multiple applications: to prevent COVID-19 in cancer and other immunocompromised patients and to halt cancer progression and treat COVID-19. This might be important for cancer patients, who could likely benefit from enhanced innate and adaptive immunity, given prior work demonstrating improved immunity mediated by OV (*28, 29, 43*).

An important concern for a vaccine is its safety. Based on the dose and the injection timeline of T-VEC, the first oncolytic virus therapy approved by FDA (*17, 44*), we provided 2 doses of OV-spike via intratumoral injection to the mice in our melanoma tumor model. No OV-spike-related adverse effects were observed after the multiple injections and no differences in body weight or temperature were observed between the saline-injected group and the OV-spike-injected group. Local infection with OV can avoid systemic toxicity. Nonetheless, we performed both i.v. and i.p. injection of the OV-spike and did not find side effects, including a substantial body temperature change. This lack of systemic side effects is consistent with our recent study that demonstrated safety of our HSV-based OV (*29*). Also, clinical studies using a similar OV have shown it is safe to treat dozens of cancer patients (*20*). Therefore, we propose that OV-spike could produce a favorable safety profile for cancer patients.

Variants of SARS-CoV-2, such as B.1.1.7, have increased transmissibility, a higher viral load, and resistance to vaccination (*45, 46*). We tested our sera from the OV-spike immunized mice on neutralization function of the B.1.1.7 variant. Compared to the titer of neutralizing WT SARS-CoV-2 infection, a similar titer of sera used to block infection by a wild-type strain could also block B.1.1.7 variant infection, which indicates that OV-spike seems to be capable of preventing infection by either the wild-type strain or the B.1.1.7 variant.

In summary, compared to existing SARS-CoV-2 vaccines which only neutralize viral infection, our vector-based OV-spike vaccine works as a dual functional agent to prevent SARS-CoV-2 infection and treat cancer progression. OV-spike shows a promising safety profile in mouse models and induces long-lasting anti-tumoral and anti-viral immune responses.

## MATERIALS AND METHODS

### Patient sample collection

Patient samples were collected from City of Hope patients and volunteers who were diagnosed with SARS-CoV-2 infection using an FDA-approved PCR-based assay. Each patient had signed a consent as part of the IRB-approved protocol 20126 at City of Hope.

### Ethics statement

All experiments using mice were conducted in compliance with federal, state, and local guidelines and with approval from the City of Hope Animal Care and Use Committee.

### Cells

The Vero *cercopithecus aethiops*-derived kidney epithelial cell line, B16 *mus musculus*-derived skin melanoma cell line, MC38 *mus musculus*-derived colon adenocarcinoma cell line, and ID8 mouse ovarian surface epithelial cell line were cultured with Dulbecco’s modified Eagle’s medium (DMEM) with 10% FBS, penicillin (100 U/ml), and streptomycin (100 μg/ml). The ID8 cells were modified to express a fly luciferase (FFL) gene (ID8-FFL) and used for *in vivo* imaging. All cell lines were routinely tested for the absence of mycoplasma using the MycoAlert Plus Mycoplasma Detection Kit from Lonza (Walkersville, MD).

### Generation of OV-spike

OV-spike was generated using the fHsvQuik-1 system, as previously described(*29, 30*). The full-length SARS-CoV-2 S protein was fused with the oHSV glycoprotein D transmembrane domain and intracellular domain. The fusion protein was inserted into pT-oriSIE4/5 following the HSV pIE4/5 promoter to construct pT-oriSIE4/5-spike. pT-oriSIE4/5-spike or pT-oriSIE4/5 was recombined with fHsvQuik-1 for engineering OV-spike and OV-Q1, respectively. Vero cells were used for propagating and titrating the viruses. Virus titration was performed using plaque assays. Briefly, monolayer Vero cells were seeded in a 96-well plate. After 12 hours, these cells were infected with gradient-diluted viral solutions. The infection media were replaced with DMEM supplemented with 10% FBS at 2 hours post infection. GFP-positive plaques were observed and counted with a Zeiss fluorescence microscope (AXIO observer 7) 2 days after infection to calculate the viral titer. To concentrate and purify the OV-Q1 and OV-spike viral particles, the culture media containing viruses were harvested and centrifuged at 3,000 x g for 30 minutes. Then the supernatants were collected and ultra-centrifuged at 100,000 x g for 1 hour. The pellets of virus were resuspended with saline as needed.

### SARS-CoV-2 neutralization, cell infection, plaque assay, and immunoplaque assay

The following reagent was obtained through BEI Resources, NIAID, NIH: SARS-Related Coronavirus 2, Isolate USA-WA1/2020, NR-52281 and SARS-Related Coronavirus 2, Isolate USA/CA_CDC_5574/2020, NR-54011. Virus isolates were passaged in Vero E6 cells (ATCC CRL-1586) as previously described (*47*). Virus concentration was determined using immunoplaque assay (also called focus forming assay) (*48*) or plaque assay. For the plaque assay, 120 pfu SARS-CoV-2 was incubated with diluted sera for 2 hours at 37 °C. Then Vero E6 cells were infected with 250 µl virus-sera mixture for 1 hour. After infection, the medium containing virus was removed, and overlay medium containing FBS-free DMEM and 2% low-melting point agarose was added. At 72 hours post infection, infected cells were fixed by 4% paraformaldehyde (PFA) overnight, and stained with 0.2% crystal violet. For the immunoplaque assay, 100 pfu of live SARS-CoV-2 were incubated with diluted sera for 2 hour and then the virus antibody mixture was added to Vero E6 cells for 1 hour at 37 °C. After 1 hour the virus containing medium was removed, overlayed with medium containing methylcellulose and 2% FBS DMEM, and incubated at 37 °C. At 24 hours after infection, infected cells were fixed by 4% paraformaldehyde for 20 minutes at room temperature and then permeabilized by 0.52% Triton X-100/ PBS solution for 210 minutes at room temperature. SARS-CoV-2 viral nucleocapsid protein (NP) was detected using the anti-NP protein antibody (Cat. # PA5-81794, Thermo Fisher) diluted 1:10000 in 0.1% tween-20/1%BSA/PBS solution as a primary antibody, followed by detecting with an anti-rabbit secondary antibody (Cat. # ab6721, Abcam) at a 1:20,000 dilution. Plates were washed three times between antibody solutions using 0.5% tween-20 in PBS. The plates were developed using 3,3′,5,5′ Tetramethylbenzidine (TMB) and then scanned using Immunospot S6 Sentry (C.T.L Analyzers). Neutralization titers for the immunoplaque assay are defined as a 50% reduction in plaque forming units relative to the untreated wells.

### Negative staining electron microscopy

OV-Q1 and OV-spike virus specimens at certain concentrations were absorbed to glow-discharged, carbon-coated 200 mesh Formvar grids. Samples were prepared by conventional negative staining with 1% (w/v) uranyl acetate. Electron microscopy images were taken on a FEI Tecnai 12 transmission electron microscope equipped with a Gatan OneView CMOS camera.

### Immuno-electron microscopy

For immunogold labeling, 5 µl of virus suspension was absorbed to glow-discharged carbon coated Formvar grids for 2 minutes. After rinsing in PBS containing 0.05% bovine serum albumin, the grids were incubated with mouse anti-spike antibody (Cat. # 40591-MM43, SinoBiological) at 1/500 dilution for 15 minutes. After washing, the grids were incubated with a 10 nm gold particle-conjugated goat anti-mouse IgG(H+L) (Cat # EM. GMHL10 BBI Solutions) at 1/50 dilution for 15 minutes. Finally, the immunolabeled samples were negatively stained with 1% (w/v) uranyl acetate for 10 seconds. The electron microscopy images were taken on a FEI Tecnai 12 transmission electron microscope equipped with a Gatan OneView CMOS camera.

### Western blot

Concentrated samples of OV-Q1 and OV-spike were mixed with NuPAGE™ Sample Reducing Agent (Cat. # NP0008, Thermo Fisher Scientific). The samples were heated at 70°C for 10 minutes and then loaded on 15% SDS-PAGE gel. The proteins were transferred onto polyvinylidene difluoride (PVDF) membrane (Minipore), and the membrane was blocked with 5% milk in PBST for 1h at room temperature (RT). Mouse anti-spike antibody was diluted at 1:1000 in PBST containing 1% BSA and incubated with the membrane at 4°C overnight. The membrane was then washed with PBST on a shaker 3 times and incubated with HRP-conjugated goat anti-mouse IgG diluted as 1:2,000 for 1 hour at RT. Pierce™ ECL Western Blotting Substrate (Cat. # 32209, Thermo Fisher Scientific) was added to the membrane, and the blots were imaged by FluorChem E (ProteinSimple).

### Quantitative real-time PCR

OV-Q1- and OV-spike-infected Vero cells were harvested after 48 hours post infection, and viral DNA was extracted from the cells using the Qiagen DNeasy Kit (Cat. # 69504). The exacted DNA was used as a quantitative real-time PCR (qPCR) template. The sequence of the spike forward primer was: 5’-TGGATTTTTGGCACCACCCT-3’ and the reverse primer was: 5’-AGACTCCCAGGAATGGGTCA-3’. The standard curve was generated using synthesized pTwist-spike as a qPCR template. The absolute copy number of OV-Q1- and OV-spike-infected Vero cells was calculated according to the standard curve.

### Binding affinity of mouse serum samples to S protein

His-tagged full-length SARS-CoV-2 S protein (50 ng) (Cat. # 40589-V08B1, Sino Biological) was used as a coating reagent. The plate (Cat #3361, Corning) was incubated with a serial dilution of mouse serum samples for 2 hours at RT. HRP-conjugated goat anti-mouse IgG antibody (Cat. #05-4220, Invitrogen) was used for detection. Absorbance was measured at 450 nm by a Multiskan™ FC Microplate Photometer (Fisher Scientific).

### VSV-SARS-CoV-2 infection

The VSV-SARS-CoV-2 chimeric virus expressing GFP was kindly provided by Sean Whelan at Washington University School of Medicine. The virus was decorated with SARS-CoV-2 S protein in place of the native glycoprotein G (*24*). Before VSV-SARS-CoV-2 infection, mouse serum samples were inactivated in a 56 °C water bath for 30 minutes, and serial dilutions were made. Vero cells (1.5-2×10^4^) were seeded 24 hours before the infection in a 96-well plate. VSV-SARS-CoV virus and the indicated amount (5 µl, 2.5 µl, 1.25 µl, 0.6 µl and 0.3 µl) of the inactivated mouse sera were preincubated at 37 °C for 2 hours and then added to the cells. The infectivity was measured by detecting GFP fluorescence using a Zeiss fluorescence microscope (AXIO observer 7) and measured as the percentage of GFP positive cells analyzed with a Fortessa X20 flow cytometer (BD Biosciences) at 24 hours post infection. VSV-SARS-CoV-2 neutralization titer was determined at the highest dilution of which GFP% is lower than control groups.

### In vivo mouse model

Six-to eight-week-old female BALB/c or C57BL/6 mice were purchased from Jackson Laboratories (Bar Harbor, Maine). OV-spike (1×10^6^ pfu or 5×10^5^ pfu) was i.v. injected on day 0 and day 14, and 1×10^6^ pfu OV-Q1 was injected as a control. For i.p. injection, 2×10^6^ pfu or 1×10^6^ pfu OV-spike was injected on day 0 and day 14, and 2×10^6^ pfu OV-Q1 was injected as a control. Peripheral blood samples were collected once a week. The body temperature of mice was monitored daily for 3 days after vaccination.

The B16 melanoma mouse model was established by injecting 5×10^5^ B16 cells s.c. 5 days before OV or saline injection into C57BL/6 mice. On day 0 and day 2, 2×10^6^ pfu OV-Q1 or OV-spike was intratumorally injected and saline was injected as a control. Tumor size was monitored every 3 days. Peripheral blood samples were collected once a week after treatment. The mice were euthanized by ketamine/xylazine at 100/10 mg/kg when the tumor volume was over 1500 mm^3^. The MC38 and ID8 colon adenocarcinoma and ovarian cancer mouse models were established by injecting 5×10^5^ MC38 cells or 1×10^6^ ID8 cells i.p. 4 days before OV or saline injection into C57BL/6 mice. On day 0, 2×10^6^ pfu OV-Q1 or OV-spike were i.p. injected and saline was injected as a control. The other 2 injections were performed on day 7 and day 14. Luciferase-based *in vivo* images were taken from 6 days after first dose of OV or saline injection to evaluate the tumor development. Peripheral blood samples were collected once a week after treatment. The mice were euthanized by ketamine/xylazine at 100/10 mg/kg when moribund and when the body weight had increased by over 20%. Experiments and handling of mice were conducted under federal, state, and local guidelines and with an approval from the City of Hope Animal Care and Use Committee.

### ELISpot

ELISpot assays for the detection of IFNγ-secreting mouse splenocytes were performed with mouse IFNγ kit (Cat. # mIFNg-1M/2, ImmunoSpot). The 96-well plate was coated with an IFNγ capture antibody at 4 °C overnight. Fresh mouse spleen cells (3×10^5^) were added to each well along with the spike peptide pool of 1.6 µg/ml. After 48 hours of incubation at 37 °C, IFNγ spots were visualized by stepwise addition of a biotinylated detection antibody, a streptavidin-enzyme conjugate and the substrate. Spots were counted using an ImmunoSpot S6 Universal Reader (CTL Europe) and analyzed using GraphPad.

### Intracellular cytokine staining and flow cytometry

Fresh mouse splenocytes were incubated with 1.6 μg/ml spike peptide pool for 24 hours at 37 °C. After treatment with brefeldin A (Cat. # 420601, Biolegend) for 4 hours, the splenocytes were stained with the extracellular markers PE-Cy ™ 7 Hamster Anti-Mouse CD3e (Cat. # 552774, BD Pharmingen), APC-Cy7 Rat Anti-Mouse CD4 (Cat. # 552051, BD Pharmingen), and CD8 alpha Monoclonal Antibody (KT15), FITC (Cat. # MA5-16759. Invitrogen) for incubation on ice for 25 minutes. The cells were washed once with PBS and fixed and permeabilized for 30 minutes avoiding direct light at RT using the fixation and permeabilization kit (Thermo Fisher Scientific) according to the manufacturer’s protocol. After washing once with permeabilization wash buffer, the cells were stained with PE Rat Anti-Mouse IFNγ (Cat. # 554412, BD Pharmingen) for 30 minutes on ice. At the same time, fresh mouse splenocytes were isolated and stained with FITC Rat Anti-Mouse CD45 (Cat. # 553080, BD Pharmingen), PE-Cy ™ 7 Hamster Anti-Mouse CD3e (Cat. # 552774, BD Pharmingen), Alexa Fluor® 700 Rat Anti-Mouse CD335 (NKp46) (Cat. # 561169, BD Pharmingen), and PE Rat anti-Mouse CD107a (Cat. # 558661, BD Pharmingen) to analyze the NK cell percentage and CD107a expression level. Flow cytometry data were acquired on a BD LSRFortessa X-20 (BD) and analyzed by FlowJo software.

### Histopathology

Mice organs including lung, brain, kidney, and liver were freshly isolated from 1×10^6^ pfu OV-spike-, OV-Q1- or saline i.v. injected BALB/c mice on day 70 post injection. Samples were placed in 10% neutral buffered formalin for a minimum of 72 hours. After paraffin embedding, 4 μm-thick sections were cut from the blocks. H&E staining were performed by the Pathology Cores, City of Hope.

### Statistical analysis

Prism software v.8 (GraphPad, CA, USA) and SAS v.9.4 (SAS Institute. NC, USA) were used to perform statistical analyses. For continuous endpoints that are normally distributed or normally distributed after logarithmic transformation, such as mean fluorescence intensity or copy number, a Student’s *t* test was used to compare the 2 independent groups. One-way ANOVA models or generalized linear models were used to compare 3 or more independent groups. For data with repeated measures from the same subject, linear mixed models were used to account for the variance and covariance structure due to repeated measures. Survival functions were estimated by the Kaplan–Meier method and compared by the two-sided log-rank test. All tests were two-sided. P-values were adjusted for multiple comparisons by Holm’s procedure. A P value of 0.05 or less was considered statistically significant.

## Supplementary Materials

Fig. S1. Intravenous injection of OV-spike induces anti-S antibody production in mouse sera.

Fig. S2. Sera collected from OV-spike vaccinated mice show strong neutralization function against VSV-SARS-CoV-2 infection.

Fig. S3. OV-spike and OV-Q1 inhibit ovarian ID8 cell tumor growth.

Fig. S4. The neutralization assay against the live wild-type and the B.1.1.7 virus strain infection of the sera from vaccinated mice bearing tumors.

Fig. S5. No significant difference of anti-S-specific antibody production between the tumor models. Fig. S6. ELISpot assay of vaccinated mice with or without tumors.

Fig. S7. Lack of side effects in OV-spike vaccinated mice.

## Acknowledgements

The authors thank Sean P.J. Whelan for sharing the VSV-SARS-CoV-2 virus. We also thank Veronica Karpinski for her technical support on ELISpot assay.

## Author contributions

J. Yu, M.A. Caligiuri, and L. Tian, conceived, designed, and supervised the project. Y. Sun, W. Dong, L. Tian, Y. Rao, C. Qin, S. Ma, K. Yu, B. Xu, J. Zhuo, R. Ma, S. Jaramillo, E. Settles, P. Keim, Z. Li, S. Dadwal, B. Barker, and P. Feng designed and/or conducted experiments. Y. Sun, L. Tian, J. Zhang performed data analyses. Y. Sun, L. Tian, J. Yu, and M.A. Caligiuri wrote, reviewed and/or revised the paper. All authors discussed the results and commented on the manuscript.

## Data and materials availability

All data associated with this study are present in the paper or the Supplementary Figures.

