## supplemental data for "Dual roles of a novel oncolytic viral vector-based SARS-CoV-2 vaccine: preventing COVID-19 and treating tumor progression"

**Supplementary Figures and legends**

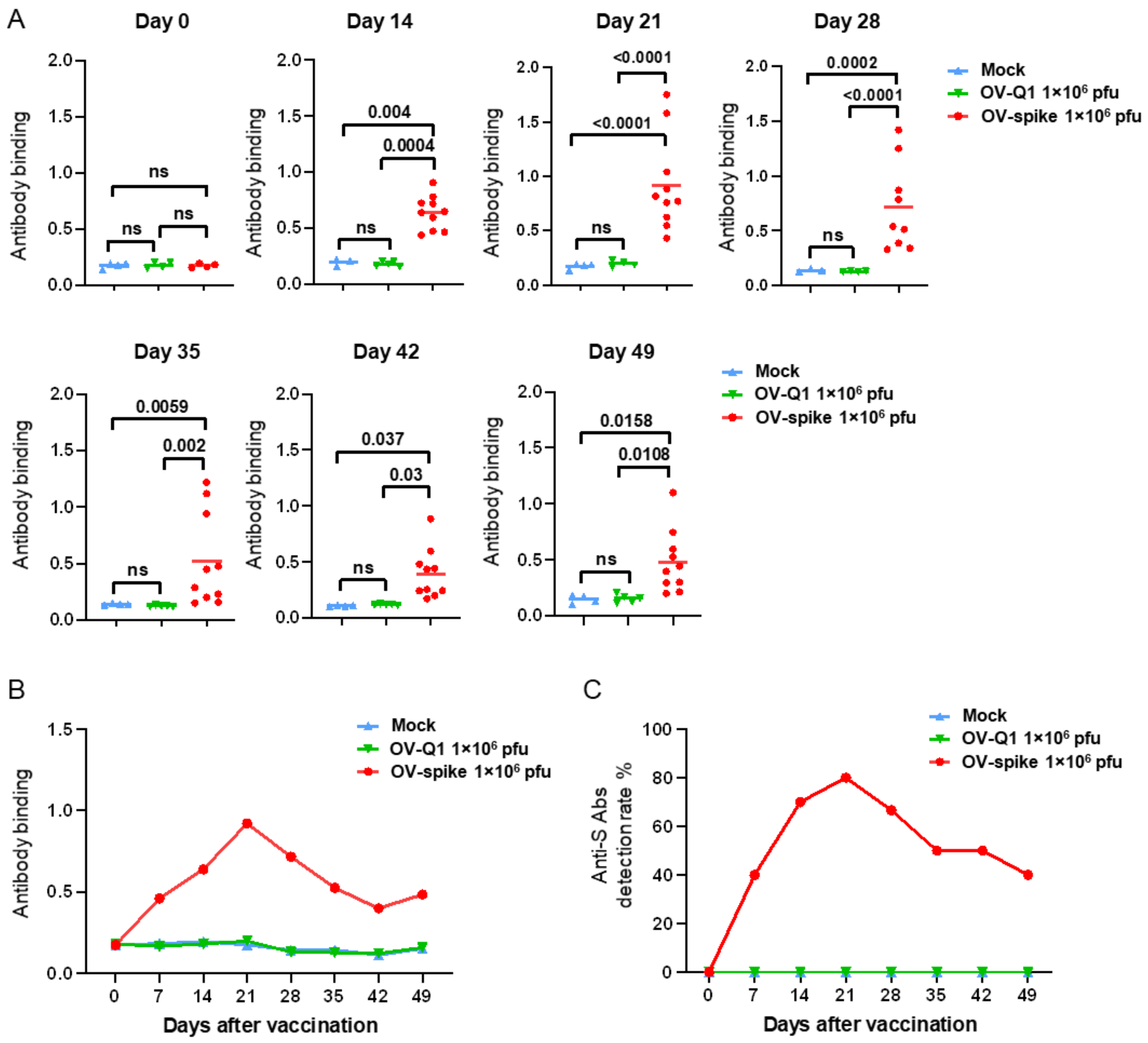


**Fig. S1. Intravenous injection of OV-spike induces anti-S antibody production in mouse sera.** C57BL/6 mice were vaccinated by i.v. injection with $1\times10^6$ pfu OV-spike, $1\times10^6$ pfu OV-Q1 or saline on days 0 and 14. **(A)** Anti-S antibody production of the sera from these mice was assessed on days 0, 21, 28, 35, 42, and 49 by a S protein-based ELISA. **(B)** The overview of the immune responses of the sera from these mice was assessed from days 0 to 49 by an S protein-based ELISA. **(C)** Anti-S production rates in vaccinated mice at indicated days. Data in A are shown in mean value, and statistical analyses were performed by one-way ANOVA with P values corrected for multiple comparisons by Bonferroni method multiple comparisons test (n = 3 to 5 mice for mock group and OV-Q1 group, n = 4-10 mice for OV-spike group).

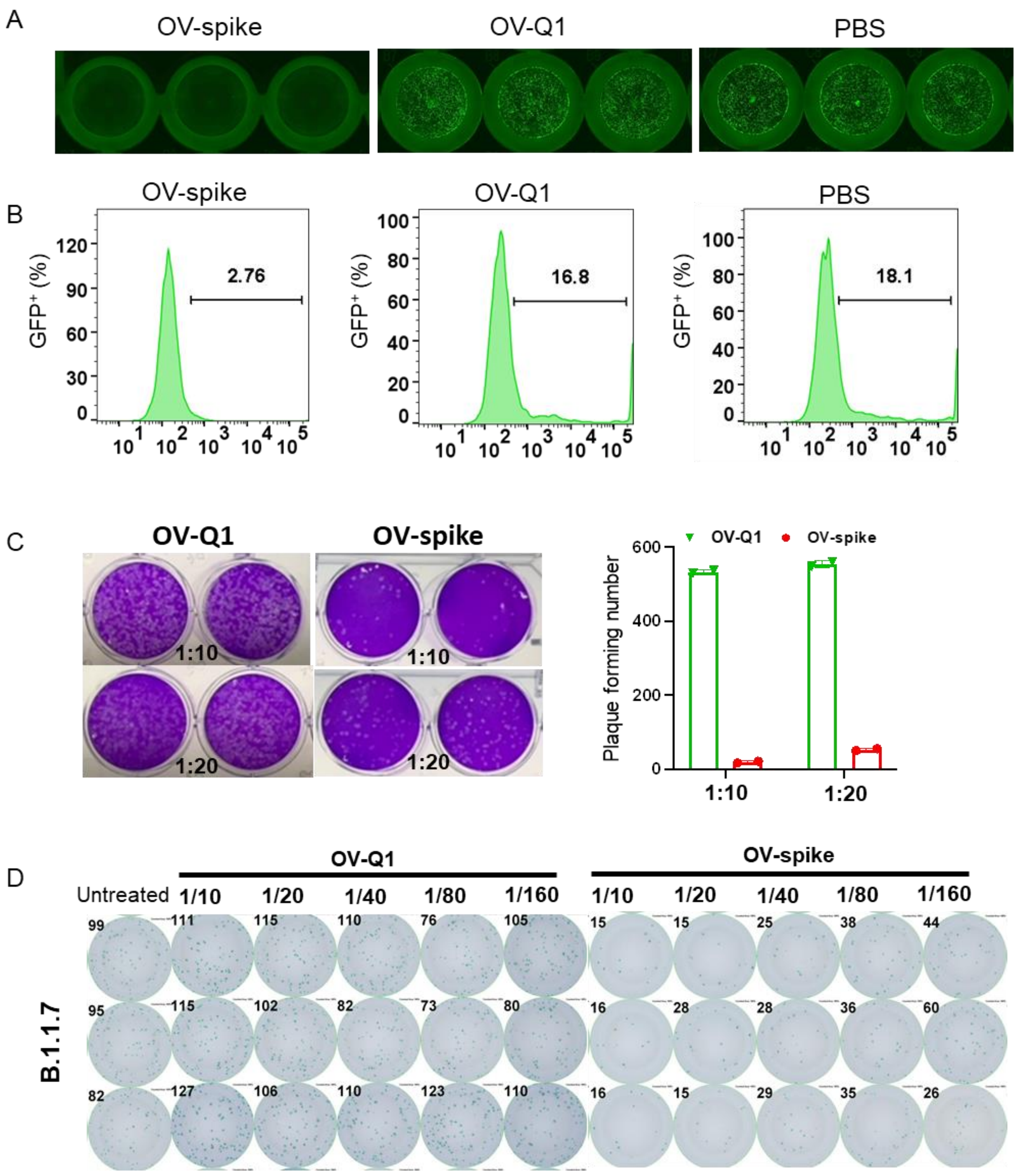


**Fig. S2. Sera collected from OV-spike vaccinated mice show strong neutralization function against VSV-SARS-CoV-2 infection. (A and B)** Cells infected with VSV-SARS-CoV-2 preincubated with different sera were then imaged at 48 hours post infection with a fluorescence microscope (A) and corresponding infectivity shown by the percentage of $GFP^+$ cells was measured by flow cytometry (B). **(C)** The left panel showed the neutralization assay against live SARS-CoV-2 infection of the sera from vaccinated mice at indicated dilution ratios. The data are summarized in the right panel. **(D)** The neutralization assay against the live B.1.1.7 variant infection of the sera from vaccinated mice at indicated dilution ratios.

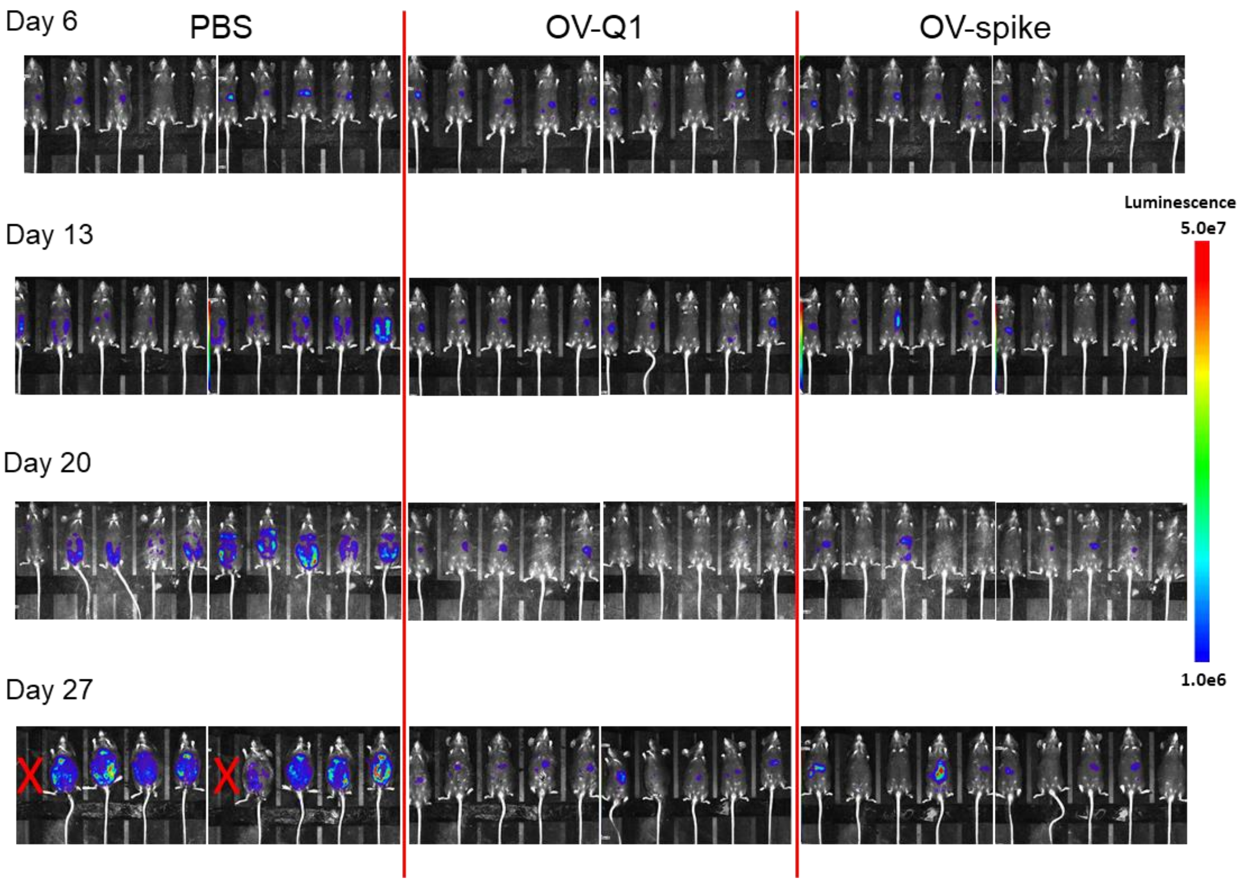


**Fig. S3. OV-spike and OV-Q1 inhibit ovarian ID8 cell tumor growth.** Time-lapse luciferase imaging of mice injected with ID8 ovarian cancer cells and vaccinated with saline(mock), OV-Q1, or OV-spike.

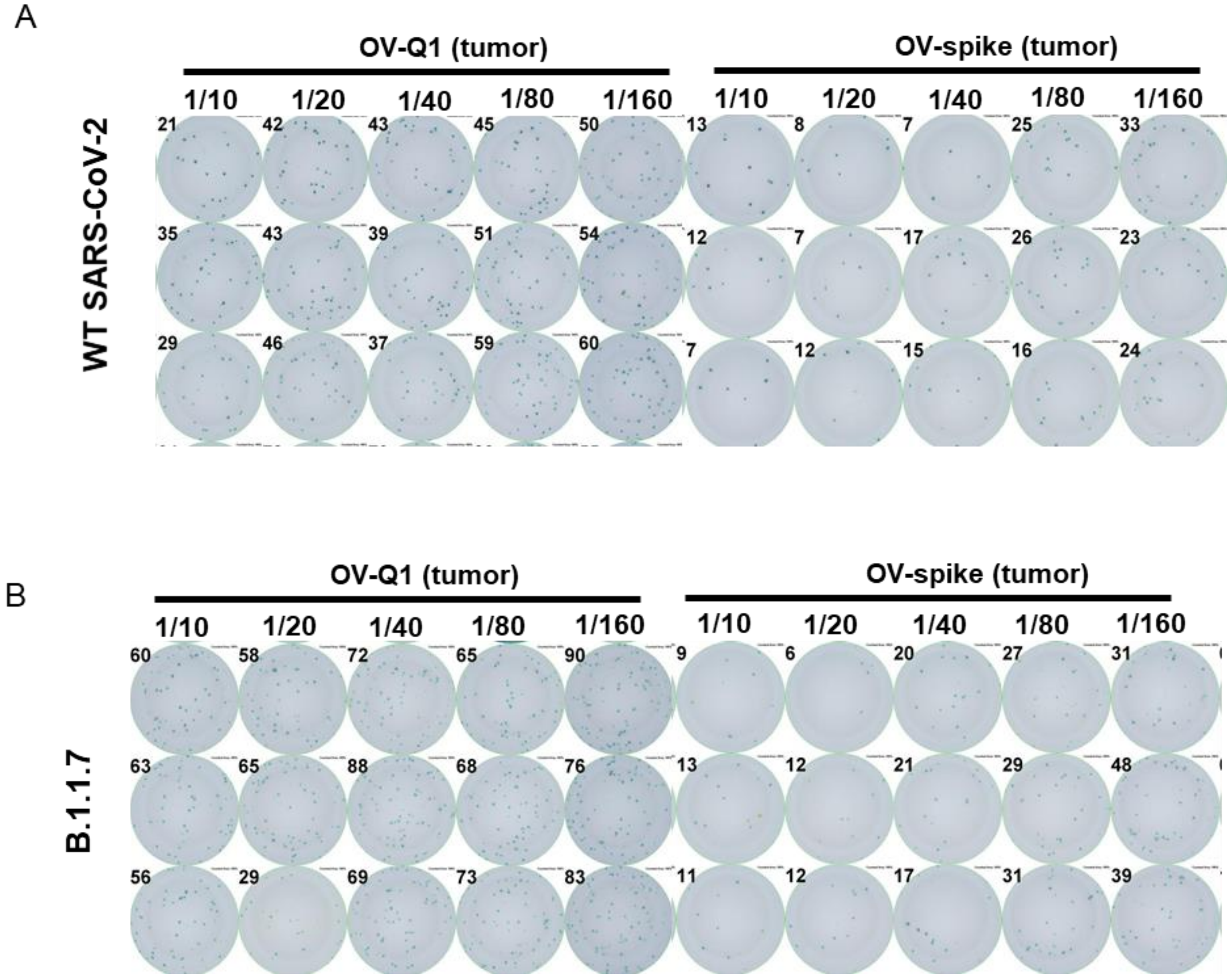


**Fig. S4. The neutralization assay against the live wild-type and the B.1.1.7 virus strain infection of the sera from vaccinated mice bearing tumors. (A and B)** The neutralization assay against the live wild-type virus strain (WT) (A) and the B.1.1.7 variant (B) infection of the sera from vaccinated mice bearing tumors at indicated dilution ratios.

A

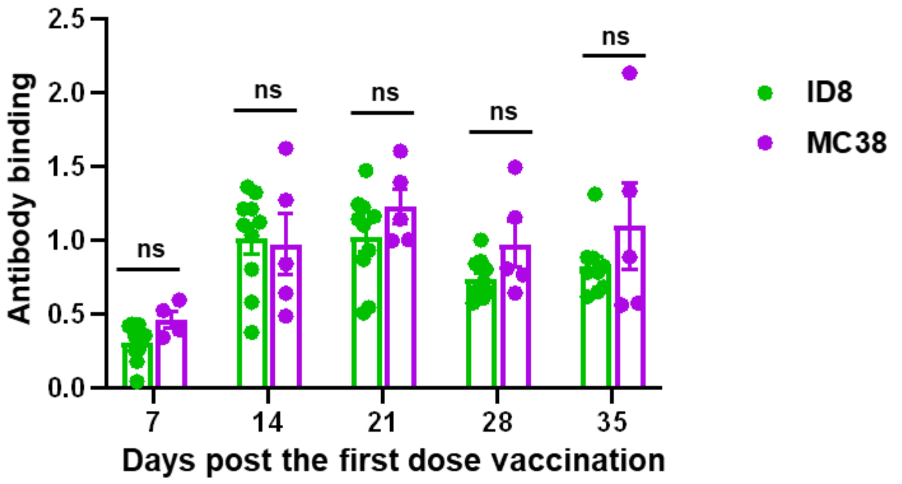


B

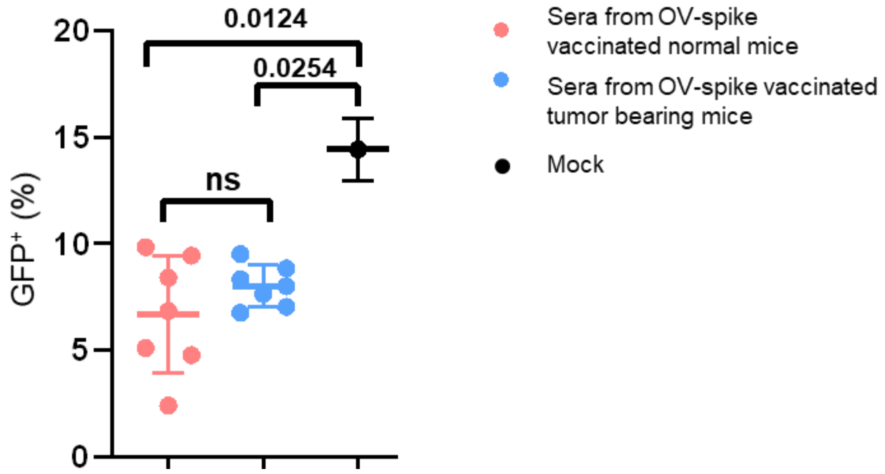


**Fig. S5. No significant difference of anti-S-specific antibody production between the tumor models. (A)** Anti-S antibody production of the sera collected from OV-spike vaccinated mice bearing ID8 tumor or MC38 tumor were assessed at day 7, 14, 21, 28, and 35 sera ELISA. **(B)** neutralization function against VSV-SARS-CoV-2 infection of sera from OV-spike-immunized mice with or without tumor. Error bars represent standard deviations. Two-sample t test with two-tail distribution was applied for (A) and one-way ANOVA was applied to compare the mean of each column with the mean of every other column with Holm-Sidak test for (B).

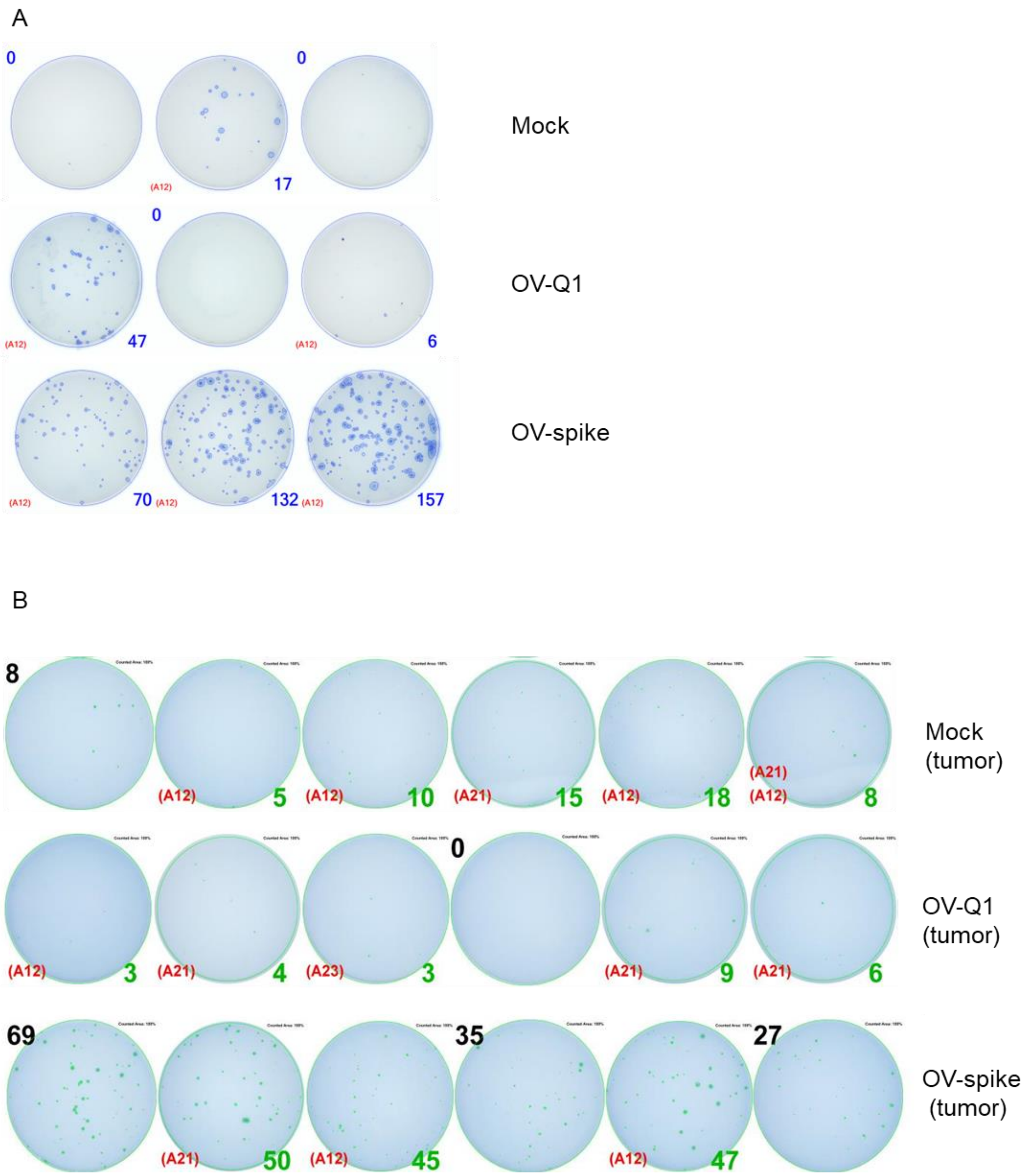


**Fig. S6. ELISpot assay of vaccinated mice with or without tumors. (A and B)** Ex vivo antigen-stimulated T cells were analyzed of sera from vaccinated mice without tumor (A) or with tumor (B). The number next to the image of each plate well indicates the number of spots in the well.

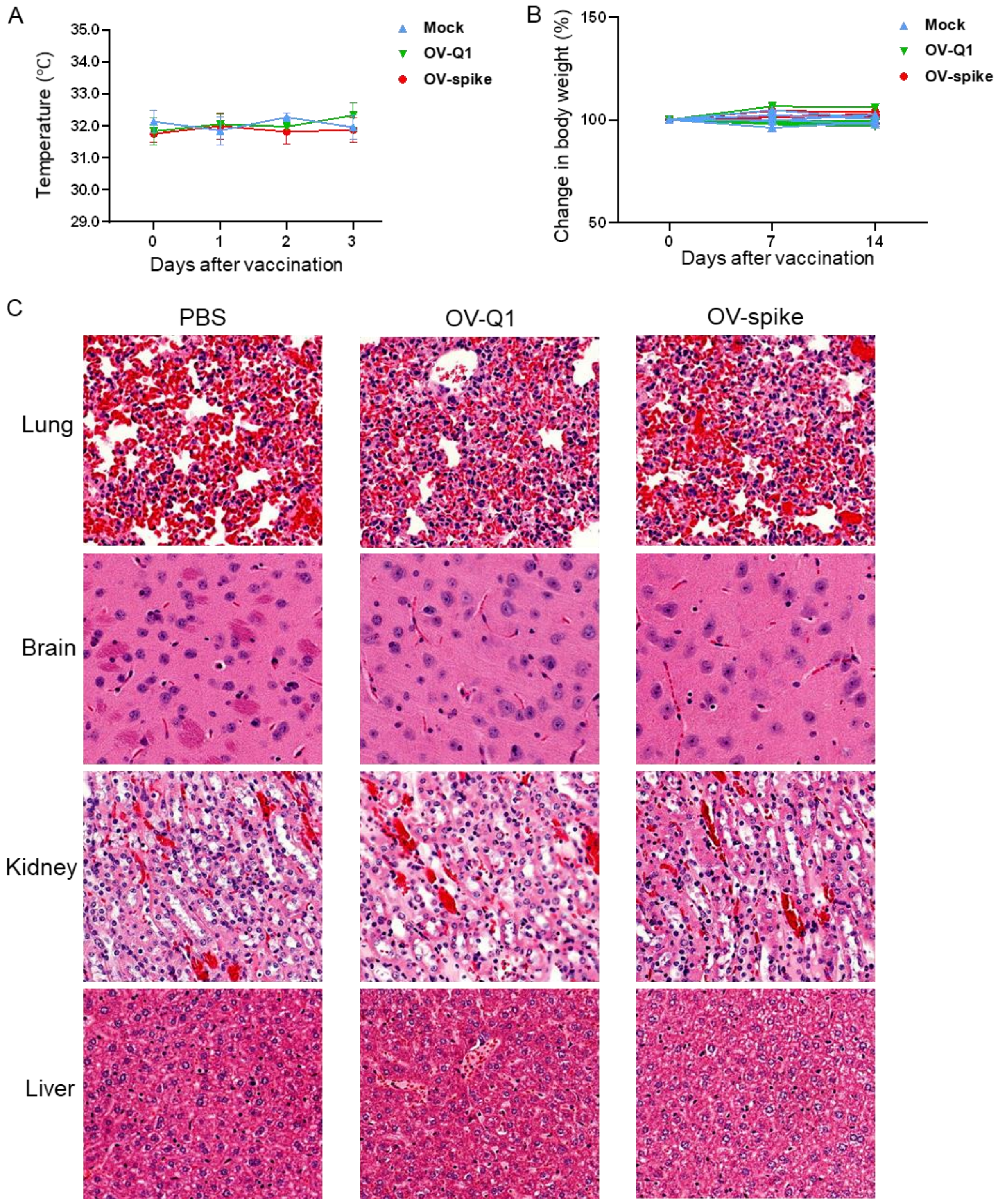


**Fig. S7. Lack of side effects in OV-spike vaccinated mice. (A and B)** Body temperature (A) and body weight (B) of mice were measured after mock, OV-Q1, or OV-spike vaccination (n = 3 per group). **(C)** H&E staining of different organs after mock, OV-Q1, or OV-spike injection.
